# Bacterial composition changes in canine plaque over periodontal disease severity and daily care practices

**DOI:** 10.1101/2023.09.13.557668

**Authors:** Ayano Watanabe, Junichi Okada, Ryo Niwa, Yukiko Inui, Kohei Ito, Yutaka Shimokawa, Miho Kihira

**Author notes:** Correspondence: Miho KIHIRA.

## Abstract

**Background:** Periodontal disease (PD) is a common oral disease in dogs and humans. Dogs have distinctly different oral environments from humans. Although common bacteria are observed in both species, profiling of the causative bacteria for the progression of PD in dogs is limited compared to humans.

**Results:** Our study examined the shifts in the bacterial community within canine plaque as PD intensifies, analyzing plaque samples from 48 dogs at various PD stages. Additionally, we examined the impact of a tooth-brushing regimen using a dental gel on twelve dogs. We revealed a correlation between the age of the dogs and the severity of PD. As PD advanced, we noted a marked increase in *Porphyromonas* abundance, a key pathogenic genus. Conversely, *Conchiformibius* prevalence diminished in higher PD levels. Furthermore, a regimen of two week brushing with a dental gel resulted in a notable decrease in *Porphyromonas* levels and five of the twelve dogs improve severity.

**Conclusions:** Our findings suggest the potential efficacy of daily brushing with dental gels, incorporating compounds proven effective in humans, for managing PD in dogs. This study demonstrates a distinct disease progression in dogs compared to humans, underscoring the need for continued research and innovation in veterinary oral healthcare.

## Background

Periodontal disease (PD) is one of the most common diseases in dogs and humans. It is estimated that more than 80% of dogs over three years of age and approximately 45% of adult humans are estimated to have PD, which can lead to tooth loss if left untreated^1–3^. PD is caused by an infection of bacteria in the oral cavity, primarily due to the accumulation of dental plaque in the cervical region, which triggers gingivitis^4, 5, 6, 7^. As PD progresses, gums can swell, bleed, and recede, forming deep periodontal. This leads to the deterioration of the gums and supporting tissues and potential tooth loss^4, 5^.

Several assessments are performed to evaluate PD stage, including probing depth, clinical attachment level, plaque index, gingivitis index, gingival bleeding on probing, involvement of furcation lesions, tooth mobility, and radiographs according to American Veterinary Dental Association criteria (AVDC)^4, 8, 9^. Anesthesia is required for ensuring accurate measurements of PD stages and minimize discomfort of dogs. Early detection and scaling is effective in preventing PD progression^10,11^. However many cases of PD are only addressed once they reached a severe stage^12^. This could be attributed to owners’ difficulty in recognizing the progression of PD and accepting the mortality risk of anesthesia for dental checkup^13, 14^. These facts highlight the necessity of improving daily care to inhibit the progression of PD.

While previous studies have extensively analyzed the prevalence, severity, and risk factors for canine PD, including poor oral care, diet, behavior, environment, and genetics^15^, the causative pathogen is still debated^16–19^. In contrast, the human oral microbiome has been extensively profiled^20, 21^. In humans, the etiology of periodontal disease is well understood through the socransky’s periodontal disease pyramid. The development of plaque biofilms is recognized as a complex process that involving the sequential participation of various bacterial complexes^5, 22^. The red complex comprises PD-associated bacteria, including *Tannerella forsythia*, *Treponema denticola*, and *Porphyromonas gingivalis*^5, 7^. Orange complex, associated with dental plaque, includes gram-negative bacteria such as *Campylobacter gracilis*, *C. rectus*, *C. showae*, *Eubacterium nodatum*, *Fusobacterium nucleatum*, *F. periodonticum*, *Peptostreptococcus micros*, *Prevotella intermedia*, *Prevotella nigrescens,* and *Streptococcus contellatus*^5^. The green, yellow, and purple complex are bacteria associated with periodontal health. The purple complex bacteria consist of *Veillonella* sp., *Actinobacillus* sp. and *Actinomyces* spp.; the yellow complex bacteria consist of *Streptococcus* spp.; the green complex bacteria consist of *Capnocytophaga* spp., *Actinobacillus actinomycetemcomitans,* and *Eikenella corrodens*^5^. The green, yellow and purple complex are colonized prior to the proliferation of the orange and red complexes, which contain gram-negative pathogenic bacteria^5^. The red complexes are found in patients with advanced periodontitis and dental plaque, and the amount of these organisms is associated with deeper pocket depth and bleeding on probing^23^. Bacteria belonging to the red complex are not observed in the absence of the orange complex. Moreover, red complex colonization increases as orange complex colonies increase. These phenomenon implies interspecific interactions, coaggregation, and metabolic interdependence^5^.

The association between PD and oral bacteria is well understood in humans, but there are also emerging findings in the canine oral microbiome^5, 24^. The oral microbiomes of dogs and humans differ greatly, and only 4.8% of shared taxa were found when comparing the oral microbiomes of canines and their owners^25^. Another study suggested that only 16.4% of the shared taxa were identified based on 16S rRNA sequence analysis^26^. Furthermore, dogs have a unique core oral microbiome^27^, and the alkaline oral environment of dogs promotes the formation of dental calculus, unlike humans^28, 29^. As the severity periodontitis in dogs increases, their oral microbiome composition changes^24, 30^. Notably, humans and dogs shared the genus of health-related bacteria, such as *Bergeyella* spp., *Morexella* spp., and *Neisseria* spp., and pathogenic bacteria, namely *Porphyromonas* spp., were observed at all PD grades in the canine oral microbiome^24, 30^. These fact indicate that bacterial species involved in the progression of PD differ from humans, even though common bacteria may be observed. However, understanding the network in these bacteria is yet limited. This raises questions about the applicability and effectiveness of bacterial tests and care methods considered beneficial in humans when applied to dogs.

In this study, we aimed to clarify the correlation between PD severity and the composition of the subgingival bacterial community in canines. As the progression of PD varies for each individual tooth, we focused on forth premolar, which is important functional tooth in dogs. We classified PD severity with previously published method^9^ and collected subgingival dental plaque under non-anesthetic conditions and profiled the bacterial compositions by 16S rRNA amplicon sequencing. PD severity was classified into three levels: Group 1 (without PD), Group 2 (mild to moderate PD), and Group 3 (severe PD), and compared each group statistically. Moreover, we sought to identify the impact of daily tooth brushing with a dental gel on PD severity and bacterial composition of the subgingival microbiome.

## Materials and methods

### Ethical considerations

This study was conducted as an observational study with banked samples. Plaque samples were routinely collected from dogs attending KINS WITH veterinary clinic (Tokyo, Japan), with owner consent obtained for the use of biological samples and clinical data for research purposes.

### Study design and Clinical evaluation of canine participants

This study comprises two cohort studies (Fig. 1). Both cohorts’ samples were collected from privately owned domestic dogs presented at the KINS WITH veterinary clinic (Tokyo, Japan) for a routine comprehensive health assessment and care between August 2022 and January 2024. In the first study (Cohort 1), cases with no treatment for periodontal disease were randomly selected from the patients of the hospital. Exclusion criteria were (1) underlying diseases, such as major systemic disease, trauma, hyperplasia, and neoplastic lesions. (2) Subjects lacking explicit consent. The study examining the effect of tooth-brushing with a gel (Cohort 2) is an independent exploratory cohort. In the second study (Cohort 2), we randomly selected dogs that participated in a dog tooth-brushing lecture held by the veterinary clinic. In addition to the inclusion and exclusion criteria of Cohort 1, cases were excluded from the study if they had (3) Failure to correctly continue veterinary-guided brushing care for two weeks in Cohort 2. No restrictions were imposed on age, breed, weight, and sex in both cohorts. These assessments were performed by a single veterinarian to minimize the variation. Age, sex, breed, plaque score, calculus score, and gingivitis score were recorded. Plaque, supragingival calculus, and gingivitis were scored from 0 to 4 at the fourth premolar following the methodology described by Perazzi^9^. Briefly, each dog was categorized into one of three groups based on the average score of plaque, calculus and gingivitis parameters: Group 1 included dogs with no PD; Group 2 dogs with mild to moderate PD; and Group 3 dogs with severe PD. In Cohort 1, forty-eight dogs were eligible (Supplementary Table 1). In Cohort 2, twelve dogs were selected for the analysis.

**Figure 1.**
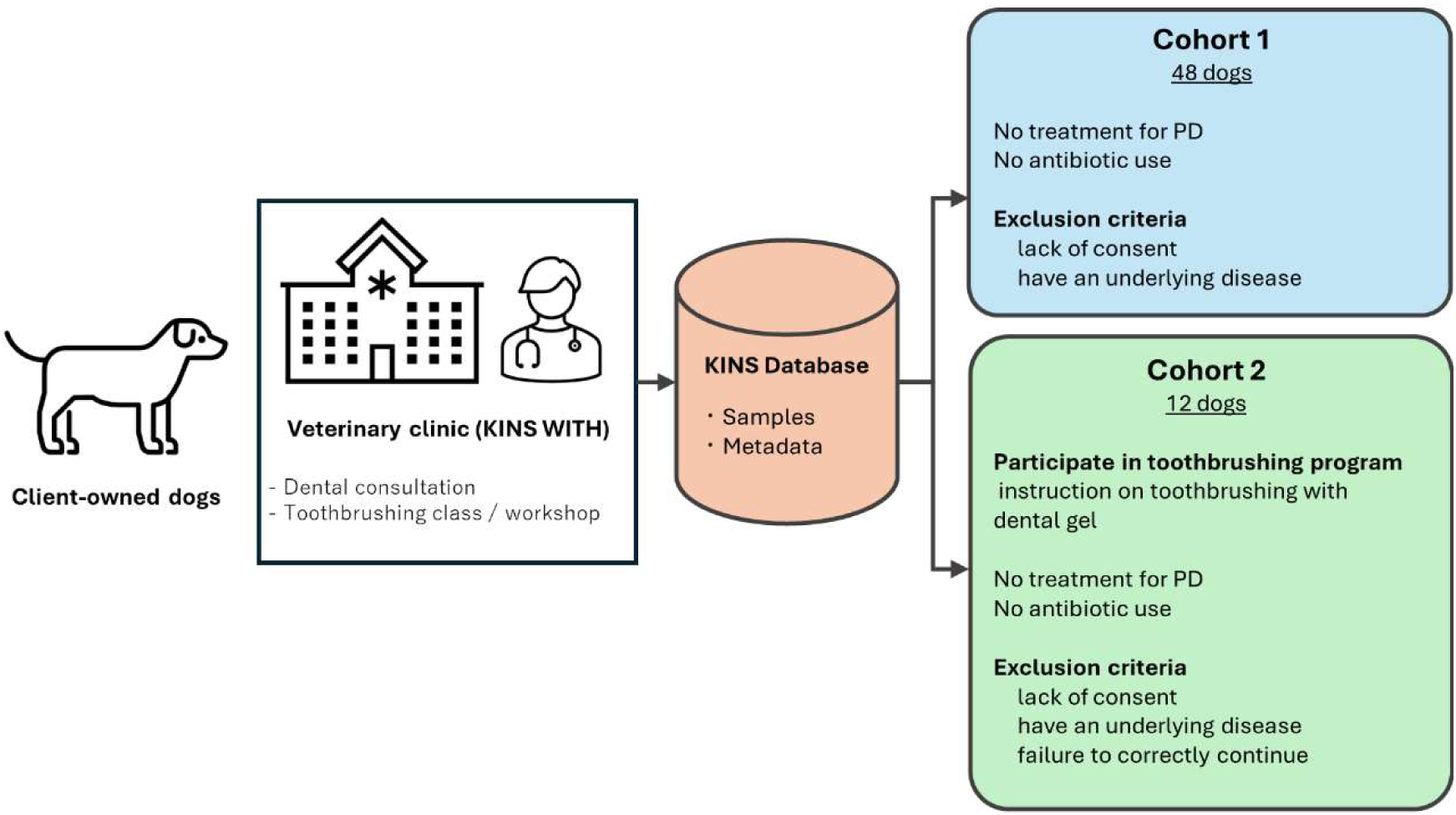
Schematic of two cohort studies.

### Veterinarian instruction Details

The veterinarian hospital provides dental examinations and instructions on teeth brushing with a dental gel (KINS, Meguro, Tokyo, Japan) as a standard practice for dogs. The veterinarian instructed owners to brush their dogs’ teeth with approximately 1 g of the dental gel using a toothbrush, at least once a day. The dental gel used in this study contained erythritol, *Chlorella* extract, *Lactobacillus crispatus*, and *L. reuteri*. The veterinarian checked up the participated dogs again two weeks after the initial sample collection (Fig. 2).

**Figure 2.**
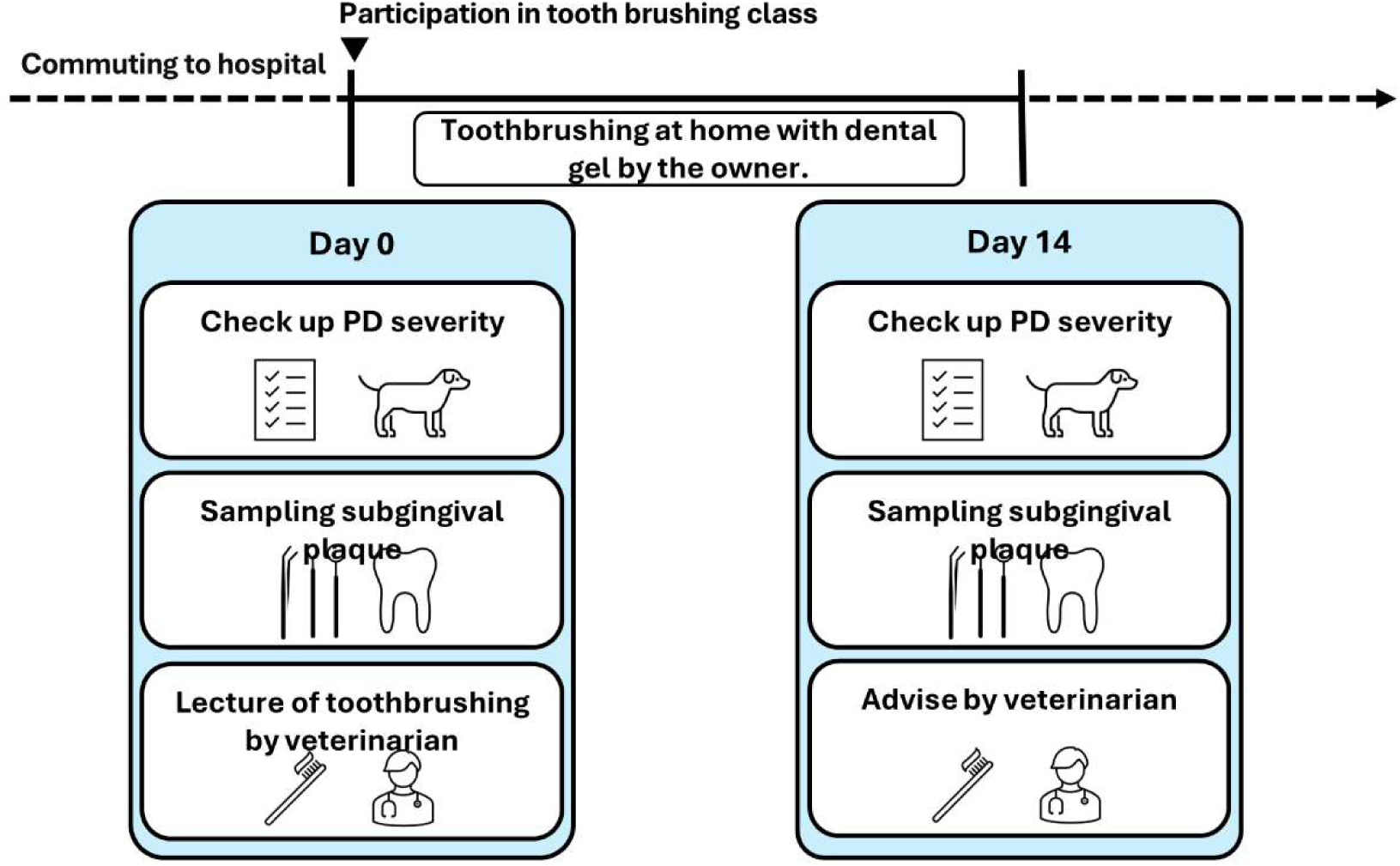
Schematic of the tooth brushing program.

### Sampling of the dental plaque microbiome

Plaque samples were collected noninvasively by using a micro applicator (COTISEN, Huanghua, Hebei, China). The target area for sampling was the fourth premolar gingival sulcus from the right side. The plaque sampling was performed by inserting the micro applicator and swiping it through a gingival sulcus. These samples were defined as subgingival plaque. After collecting the samples, we placed them in 2-mL microtubes (Eppendorf, Barkhausenweg, Hamburg, Germany) containing 250 µL of GTC solution^7^ (100 mM Tris-HCl; pH 9.0, 40 mM EDTA, and 4 M guanidine thiocyanate), temporarily stored them at 4 °C and kept them at −80 °C until DNA extraction.

### DNA extraction and PCR amplification of the 16S rRNA gene

We used the QIamp DNA Mini Kit (QIAGEN, Hilden, Nordrhein-Westfalen, Germany) according to the manufacturer’s instructions, using 100 µL of plaque samples. The hypervariable V1-V2 region of the 16S rRNA-encoding gene in DNA was amplified by PCR using the barcoded primers 27Fmod (5′- ACACTCTTTCCCTACACGACGCTCTTCCGATCT-NNNNN-AGRGTTTGATYMTGGCTCAG-3′) and 338R (5′- GTGACTGGAGTTCAGACGTGTGCTCTTCCGATCT-NNNNN-TGCTGCCTCCCGTAGGAGT-3′)^67^. We used 10×Ex Taq Buffer, 25 mM dNTP Mixture, 5 U/µL Ex Taq polymerase (Takara Bio, Kusatsu, Shiga, Japan), 20 mg/mL BSA (Roche Molecular Diagnostics, Pleasanton, CA, USA), and forward and reverse primers as the PCR mixture. PCR cycling conditions were as follows: initial denaturation at 95 °C for 2 min; 30 cycles of 95 °C for 30 sec., 55 °C for 45 sec., and 72 °C for one min., after which a final extension was conducted at 72 °C for 10 min, and the amplicon was stored at 4 °C.

### Library preparation and sequencing

The PCR amplicon was purified by AMPure XP beads (Beckman Coulter Inc., Brea, CA, USA). Purified products were quantified using the Quant-iT™ PicoGreen® dsDNA Assay Kit (Invitrogen, Carlsbad, CA, USA). Samples were treated using the MinElute PCR Purification Kit (QIAGEN, Hilden, Nordrhein-Westfalen, Germany). The products were quantified using a Qubit dsDNA BR Assay Kit (Invitrogen, Carlsbad, CA, USA) and a Kapa Library Quantification Kit (Illumina, Inc., San Diego, CA, USA), and a quality check of the samples was conducted using the 4150 TapeStation system (Agilent Technologies, Inc., Palo Alto, CA, USA). The barcoded DNA libraries were sequenced using Illumina MiSeq (Illumina, Inc., San Diego, CA, USA) with 2 × 250 bp paired end reads.

### Bioinformatics and statistical analysis

Sequence data (FASTQ) was analyzed using the QIIME2 pipeline (https://qiime2.org/, version 2022.11)^68^. Amplicon denoising and merging were conducted using the QIIME2 plugin DADA2^69^. The options TRUNC_LEN_F=198, TRUNC_LEN_R=173, FORWARDPRIMERLENGTH=20, and REVERSEPRIMERLENGTH=19 were used for the 312 bp amplicon. ASVs were then taxonomically classified using the SILVA database SSU 138^70^ and the feature classifier classify-sklearn^58^.

We constructed a phylogenetic tree and performed diversity analysis using the diversity plugin. We calculated the α-diversity and applied the Kruskal–Wallis test and pairwise Mann–Whitney test using the alpha-group significance plugin. We used the Benjamini–Hochberg method for multiple testing correction, where an adjusted *p*-value of 0.05 or less was considered statistically significant. We used the beta-group-significance plugin to determine β-diversity by PERMANOVA. A *p*-value of 0.05 or less was considered statistically significant.

We used microbiomeMarker v1.0.2 for LEfSe analysis, and only bacterial genera that reached a threshold LDA score of 4.0 with a *p*-value lower than 0.01 are shown in this study^71, 72^. We performed Tukey’s honest significant difference (HSD) test using the run_posthoc_test function of microbiomeMarker as a post hoc test for each genus whose variation was verified by LEfSe. Data visualization was conducted with R version 4.2.1, ggplot2 package^73^, ggprism^74^, and Phyloseq v. 1.40.0^75^. We used ANOVA and then further detailed the differences with Tukey’s HSD method to see if there was a correlation between the age of the dogs and their PD group. We compared the abundances of *Porphylomonas* in the two groups using the Wilcoxon signed-rank test. We considered the results significant if the *p*-value was less than 0.05.

## Results

### PD stage correlates with age

We classified the participants into three PD severity groups, utilizing criteria established in a previous study^9^. The age of the dogs in this study ranged from as young as a few months to 12 years (mean: 4.3 years, median: 4 years; Supplementary Table 1). Seven dogs were classified into Group 1, 24 dogs into Group 2, and 15 dogs into Group 3. Additionally, data on the dogs’ age and other relevant variables were collected using a questionnaire (Supplementary Table 1). The mean age of dogs differed significantly among the PD severity groups (Fig.3A). Dogs in Group 1 had a mean age of 2.86 years (SD 2.91), while those in Group2 and Group3 had mean ages of 3.52 years (SD 2.69) and 7.12 years (SD 2.94), respectively. Post-hoc comparisons revealed that the mean age of dogs in Groups 2 and 3 was significantly higher than in Group 1 (Fig. 3A; *p* = 0.005, and *p* = 0.001, respectively).

**Figure 3.**
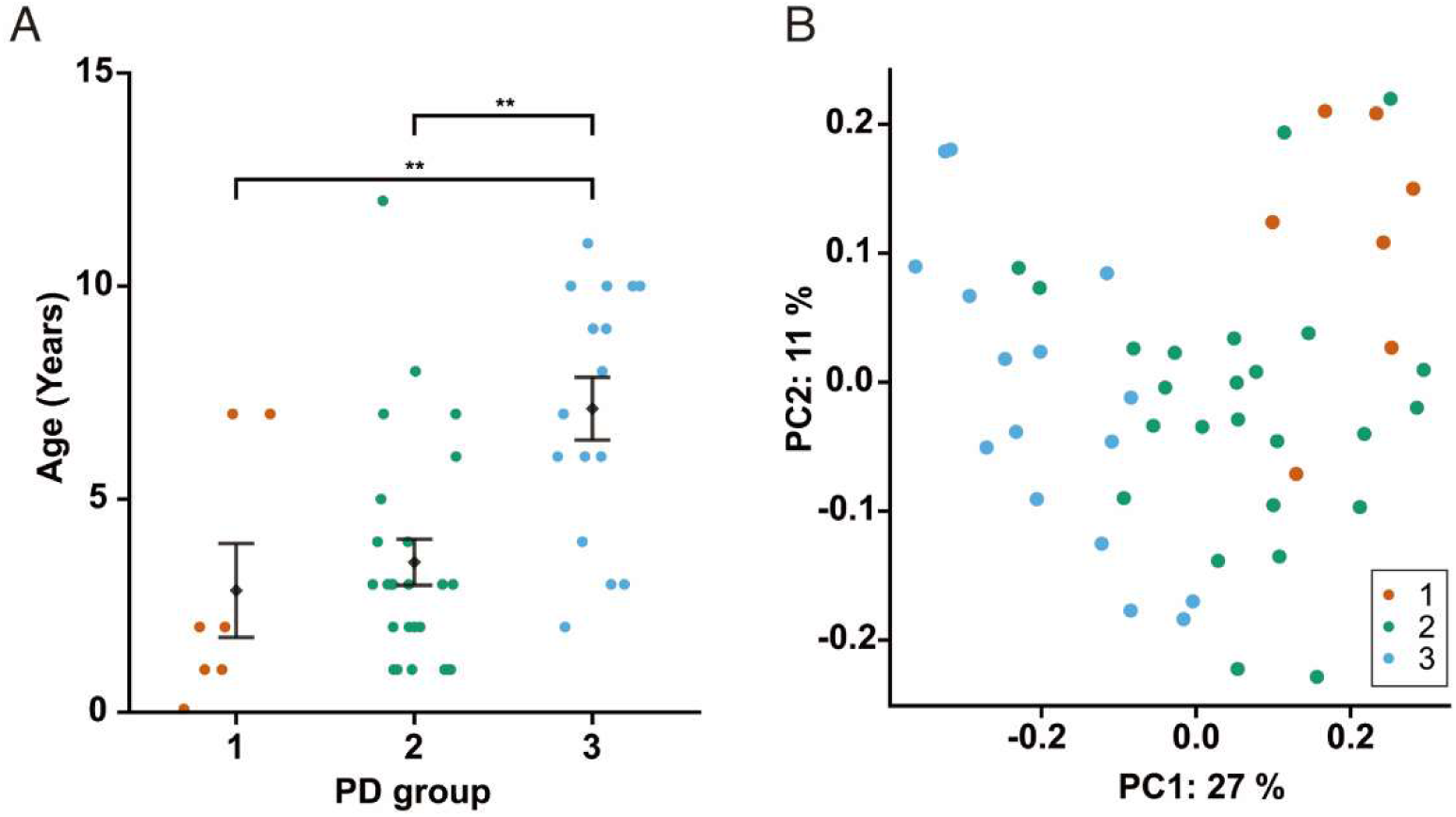
Profiling of PD grade and oral microbiome in dogs. (A) Dot plot illustrating the correlation between age and periodontal disease group. Seven, 25, and 16 dogs were classified into Group 1, Group 2, and Group 3, respectively. Statistical tests were conducted for each group and age, and <*p* = 0.01 was expressed as **. (B) Unweighted UniFrac distances for beta diversity analysis.

### Microbial structure is different among PD groups

After 16S rRNA amplicon sequencing, we removed reads classified as unassigned, mitochondrial, and chloroplast. This resulted in a minimum of 12,375 reads, a maximum of 75,038 reads, and a median of 46,934 reads across 48 samples. An alpha diversity rarefaction curve was constructed to verify that a sequencing depth of 12,375 reads adequately captured the microbial community profiles within the samples. Consequently, this sequencing depth at 12,375 was used as a rarefaction parameter in the diversity analysis (Fig. S1). For alpha diversity analysis, we calculated the Shannon diversity and observed features. Kruskal–Wallis tests adjusted for multiple comparisons revealed a significant increase in the Shannon diversity index between Groups 1 and 2 (*p* = 0.032) and Groups 1 and 3 (*p* = 0.039), with a notable rise in observed features between Groups 1 and 3 (Fig. S2A, S2B; *p* = 0.041). We used unweighted UniFrac distances for beta diversity analysis to plot the principal coordinate analysis (PCoA). The unweighted UniFrac plot distinctly separated the samples based on each PD stage. Along the first principal coordinate (PC1) axis, Group 1 samples clustered on the extreme right, Group 2 samples centered in the middle, and Group 3 samples positioned on the left. (Fig. 3B). Additionally, permutational multivariate analysis of variance (PERMANOVA) was applied to assess beta diversity and showed significant differences with *p*-values of 0.004 between Group 1 and Group 2 and <0.001 between Groups 2 and 3.

### Characteristic taxa are present in each PD group

All groups showed the presence of *Porphyromonas* spp., *Fusobacterium* spp., and *Moraxella* spp. bacteria in the median of relative abundances at the genus level (Fig. S2C). Group 1 had more *Conchiformibius* spp. than the other groups. However, *Treponema* spp. was found more frequently in Group 2 and 3 than in Group 1. As PD worsened, *Porphylomonas* spp., *Fusobacterium* spp., and *Moraxella* spp. increased, while *Conchiformibius* spp., *Neisseria* spp., *Actinomyces* spp., and *Pasteurella* spp. decreased. This shows that the bacterial composition of subgingival plaque varies with PD severity. We performed LEfSe analysis to identify genus-level features distinguishing Groups 1, 2, and 3. Group 1 contained *Conchiformibius* spp., *Pasteurella* spp. *Pasteurellaceae_uncultured*, *Actinomyces*, *Bergeyella*, *Lautropia*, and *Euzebyaceae_uncultured*. Group 2 had *Neisseria* spp., *Capnocytophaga* spp., *Corynebacterium* spp., *Frederiksenia* spp., and *Campylobacter* spp. Group 3 featured *Porphyromonas* spp., *Treponema* spp., *Bacteroides* spp., *Fretibacterium* spp., *Acholeplasma* spp., and *Leptotrichiaceae* spp. (Fig. 4A). A heatmap depicts their relative abundance (Fig. 4B). *Conchiformibius* spp. had the highest LDA score in LEfSe analysis in Group 1 and was significantly lower in Group 3 when comparing the mean value of relative abundances (Figs. 4A, 4B, and S3A; *p* = 0.003, 0.002). *Porphyromonas* spp. had the highest LDA score and was particularly enriched in Group 3, with a significantly increased abundance in Group 3 compared with the others (Figs. 2B and S3B; *p* = 0.003, 0.043).

**Figure 4.**
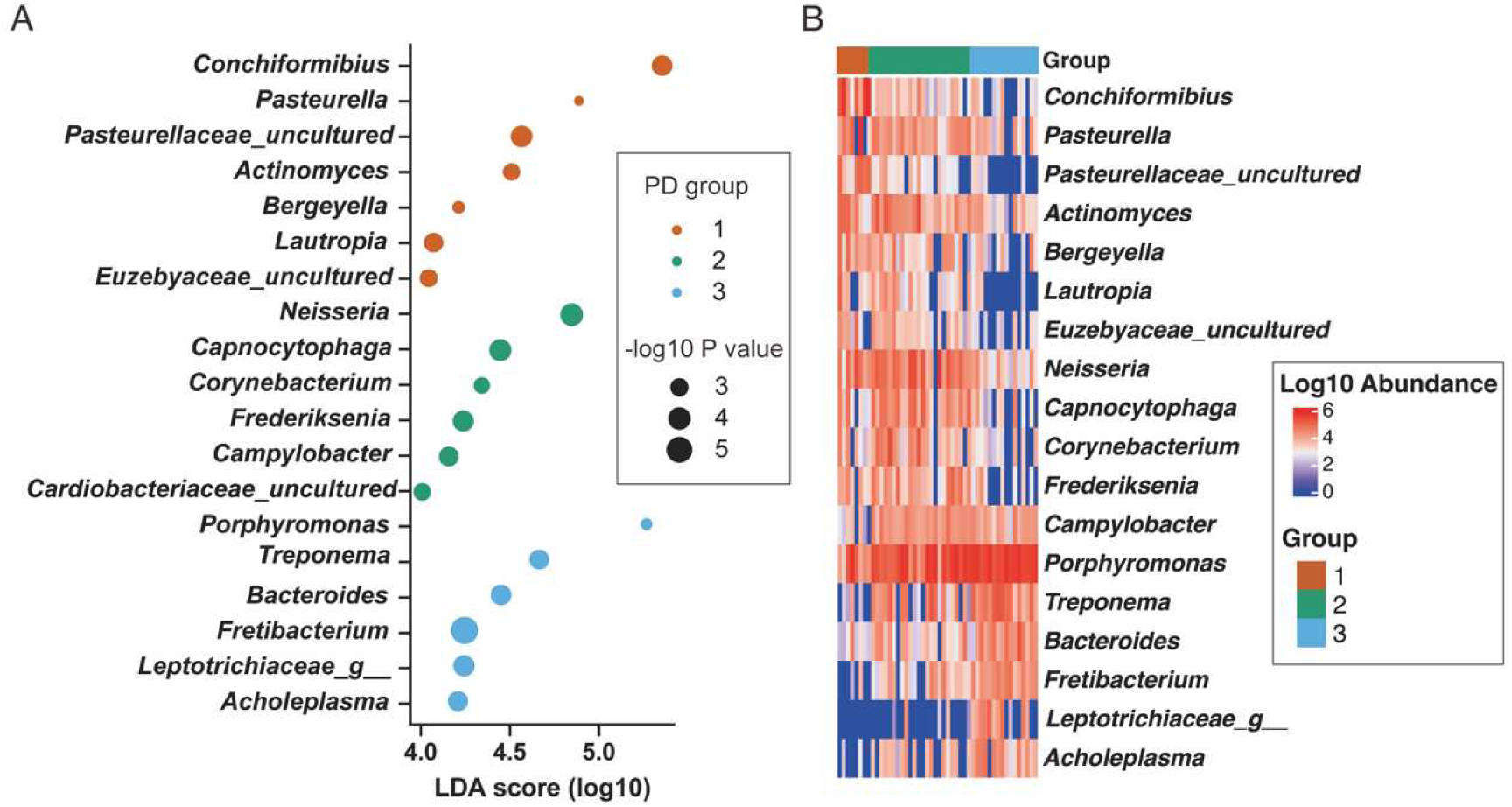
LEfSe analysis and LDA heatmap of the oral microbiome in PD and dogs. (A) Dot plot differentiating the distinct LDA values of periodontal disease groups 1, 2, and 3. (B) An LDA heatmap, visualized on a log10 scale, was formulated using post-sample normalization with the sum of values set to a million (count per million). Group 1 is represented in Red, Group 2 in green, and Group 3 in blue.

### Tooth brushing reduces the relative abundance of *Porphyromonas*

Next, we examined changes in the subgingival plaque samples before and after two weeks of tooth brushing with dental gel to verify the post effects of brushing habits on the PD-associated microbiome. We collected samples from a different group of dogs than that in our initial study and performed 16S rRNA amplicon sequencing (Supplementary Table 2). We removed reads identified as unassigned, mitochondrial, and chloroplast, resulting in a minimum of 7,942 reads, a maximum of 58,651 reads, and a median of 38,958.5 reads per sample. Before brushing, out of twelve dogs, two were in Group 1, seven were in Group 2, and three were in Group 3. After consistent brushing, two dogs moved from Group 2 to Group 1, three dogs moved from Group3 to Group 2, two remained in Group 1, five remained in Group 2, and none remained in Group 3 (Supplementary Table 2). This indicates that brushing improved PD severity in five dogs, while there was no change for the other seven (Supplementary Table 2). Upon comparing the relative abundance of bacterial genera pre- and post-two week brushing care, it was observed that *Porphyromonas spp.* decreased significantly (*p* = 0.004) in the ten dogs (Fig. 5, S5).

**Figure 5:**
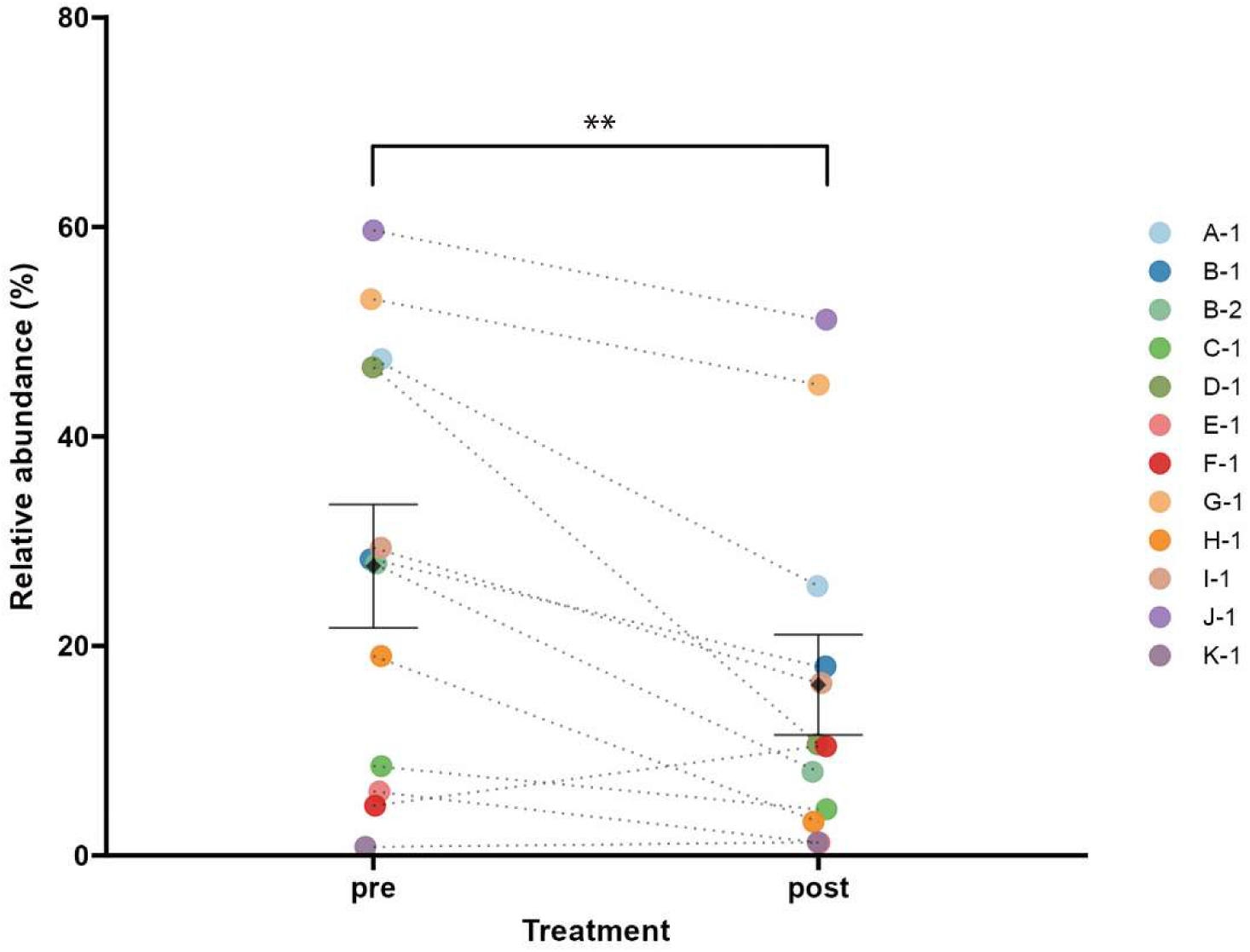
Microbiome analysis before and after brushing with gel. Changes in *Porphyromonas* pre- and post-brushing care for two weeks. The dots of each dog’s pre and post-brushing care are linked using dashed lines. Statistical tests were conducted into pre and post-brushing, and <*p* = 0.01 was expressed as **.

## Discussion

In this study, we analyzed the subgingival plaque microbiome across multiple P D stages. Our findings reveal shifts in the subgingival plaque microbiome in correlation with PD severity. Additionally, we found a correlation between age and PD severity (Fig. 3A), which is consistent with previous studies^2, 6, 31^. We also found that tooth-brushing care reduced the severity of PD dogs in five out of twelve and decreased the presence of *Porphyromonas* spp. in ten out of twelve.

In classical bacterial infections such as pyogenic infections, the diversity of the bacterial community decreases with disease progression^32^. Conversely, PD arises from a continuous process that leads to the formation of a complex microbiome formation, eventually causing periodontitis^22^. When pathogenic microbiomes are present over a long period of time, they are considered to impact the host negatively^32^.

In a previous study involving 24 dogs, 12 with gingivitis or periodontitis and 12 without PD, no significant correlation was observed in the Shannon diversity index between the two groups^33^. A separate cohort study of 587 dogs revealed a correlation between the Shannon diversity index and the average gingivitis score in medium sized dogs, but no such statistical significance was found in small or large dogs^34^. In contrast, our study, which specifically examined the fourth premolar and diagnosed PD severity, found that the subgingival plaque from dogs with PD (Groups 2 and 3) had a significant increase in the Shannon diversity index compared to the plaque of healthy dogs (Group 1; Fig. S2A, S2B). We also calculated the weighted UniFrac distance, a β-diversity evaluation weighted by sample proportions, and it showed the trajectory associated with PD severity. The observed shift in clusters corresponding to PD severity implies a change in the oral microbiome as the relative quantity of bacteria associated with PD increases. We also performed PCoA and obtained a similar conclusion from a study in 2013 that clusters were determined by PD severity using PCA^11^.Given the vast differences among dog breeds and their dietary habits and considering the varied comparisons in different studies, drawing a conclusive understanding remains challenging. Future research is crucial to provide more clarity.

*P. gingivalis* is classified as the red complex and has been extensively studied. It is a gram-negative anaerobic bacterium that is strongly associated with advanced periodontal lesions^35–37^. In dogs, the *Porphyromonas* genus is also proposed to be a potential pathogen associated with PD^11, 24, 33, 34^. We noticed that the abundance of the *Porphyromonas* spp. tended to increase with the progressive severity of PD (Fig. S2C). The LEfSe analysis identified the *Porphyromonas* spp. and *Treponema* spp. as a characteristic of severe PD (Group 3) (Fig. 4A). In addition to *P. gingivalis*, both *Tannerella forsythia* and *Treponema denticola* are also classified within the red complex for humans. Interestingly, in this study, *Treponema* spp. and *Tannerella* spp. have tend to increase with progression in PD severity (Fig. S4). Our findings suggest that *Porphyromonas* spp., *Treponema* spp., and *Tannerella* spp. may contribute to the exacerbation of canine PD, mirroring their impact in human cases. A comprehensive high-resolution analysis to allow discussion at species-level is essential for validation. Expanding our examination to include other genera in LEfSe, *Neisseria spp.*, *Capnocytophaga spp.*, and *Campylobacter spp.*, which exhibit low pathogenicity and colonize the healthy oral cavity in humans, were found to be features of Group 2 (Fig. 4A) in dogs. *Prevotella* spp. and *Fusobacterium* spp., which are classified in the orange complex in humans, were not detected as features in this analysis. *Actinomyces* spp. associated with the purple complex in humans, was found in Group 1 diminished to Group3 in dogs. Additionally, *Streptococcus* spp., belonging to the yellow category^5^, was not identified as a distinguishing feature. Given the significant differences in oral bacterial communities between dogs and humans^26^, dogs may undergo a distinct transition in the microbiome with periodontitis compared to humans. However, the bacteria implicated in the onset of severe symptoms might be similar to those in humans. It is possible that common bacteria observed in both dogs and humans can play a comparable role in the progression of PD.

Our results also highlighted *Conchiformibius* spp. as a prominent feature of Group 1 (Fig. 4A, 4B). Previous studies also clarified the presence of *Conchiformibius spp.* in healthy dogs^33, 34^ and 6-8 month old puppies^38^. However, our LEfSe analysis verified the prevalence of *Conchiformibius* spp. in Group 2, characterized by mid-moderate PD. Interestingly, its presence was minimal in Group 3 (Fig. S3A). This bacterium inhabits the oral cavity, intestines, and vagina of mammals. They are known as multicellular longitudinal division (MuLDi) species, where cells divide perpendicularly to the growth direction of growth and remain interconnected. This unique evolution is likely to suit specific conditions in the oral cavity for the bacterium^39–42^. The reduced presence of *Conchiformibius* spp. as PD worsened indicates that the genus may have difficulty adapting to the environmental alterations brought about by the progression of PD. Given that the dogs with a high presence of *Conchiformibius* spp. ranged in age from 1–7 years and only seven dogs comprised Group 1, there is interest in exploring its potential as a probiotic approach.

In dogs, several reports used oral microbiomes to monitor PD. Sampling from various niches in the oral cavity, such as maxillary arcade, supragingival plaque, subgingival plaque, and salivary was chosen previously^24, 43, 44^. While the microbiome, excluding saliva, reflect the status of PD, subgingival plaque microbiome had a higher correlation coefficient with the gingivitis score^44^. Additionally, various stages of PD can be observed in the teeth of a single dog. Therefore, we utilized forth premolar subgingival plaque samples as an indicator using micro applicator. A human study reported that microbial diversity increased as periodontal pockets deepened and different microbiome characteristics, supporting the idea that the progression of PD, leading to deeper periodontal pockets, creates new habitats where additional bacterial species can colonize^45^. In this study, we observed that microbial diversity in subgingival plaques correlated with PD severity, accompanied by changes in bacterial genus occupancy rates. This suggests that, similar to humans, the progression of PD in dogs may be associated with the emergence of new bacterial species, potentially introducing more anerobic bacteria.

Several routine oral hygiene methods have been proposed for modulate oral microbiome composition and reduce inflammation in human. Water flossing decrease amount of anerobic bacteria and alleviate inflammation compared to controls^46^. Tooth-brushing with enzyme- and protein-containing toothpaste led to changes in the bacterial community over time^47^. Several materials have been reported to manage PD symptoms in human. Erythritol reduces plaque accumulation and the presence of bacteria in both saliva and plaque^52^. *Chlorella* extract has been suggested as a potential source of physiologically active compounds with anti-tumor, anti-inflammatory, antioxidant, and antibacterial properties^54^. *Lactobacillus crispatus,* administered orally to mice infected with *P. gingivalis,* reduced periodontal symptoms. Similar effects were seen in humans, with lower plaque scores, a reddish tinge, and gingival swelling scores^55^. Ingestion of *Lactobacillus reuteri* has been reported to decrease the amount of periodontal pathogens in subgingival plaque^56, 57^. One previous study suggested that tooth brushing after scaling can effectively limit the increase in oral bacterial counts^48^. However, there are also reported that periodontal pathogen levels return to baseline after 6 months, regardless of bacterial species even with supportive therapy^49^. Periodontal pathogens are present throughout the entire oral cavity^43, 50^, allowing the oral microbiome to return over time. These finding mean that physical methods are effective in controlling oral bacterial amounts but have low selectivity among pathogenic bacteria. Notably, dog consumption of dental gum was reported improve PD symtoms^51^and the shift in bacterial genera varied accross the type of dental gum used^52^. In this study, the dental gel used in tooth brushing lecture includes several effective ingredients reported in human. Erythritol in the dental gel has been reported to have bacteriostatic effects against PD-associated bacteria in dogs^53^. Our observation showed that the use of dental gel during brushing reduced the presence of *Prohyromonas spp.*, which increased in the severe PD stage, indicating a protective effect. These findings suggest that habit of tooth brushing with application include effective ingredients can influence the pathogenic bacterial community composition.

In dogs, severe PD is irreversible, requiring prompt intervention and oral care for daily maintenance. Thus, our study, which captures shifts in the subgingival microbiome alongside the pathological development of PD, holds valuable insights. However, a potential limitation of this study is that we did not apply anesthesia for teeth observation. Consequently, this research did not include X-rays and probing, which are criteria for assessing PD stages in AVDC’s standard. These limitations restrict our capacity to investigate the relationship between alveolar bone loss, central PD pathology, and the subgingival microbiome. Incorporating more precise diagnostic methods in upcoming work could provide deeper insights into the connection between PD progression and the presence of *Porphyromonas spp..* Another limitation of the present study is that we used amplicons of the V1-V2 region of the 16S rRNA gene sequence, which limits the resolution to identify bacteria only at the genus level^58^. DNA extraction and PCR amplification also introduce various biases^59, 60^, affecting the accuracy of the relative composition of the oral microbiome. We cannot discuss whether species considered as biomarkers for human PD are also disease-causing species in dogs due to the resolution by amplicon sequencing^60^. Therefore, it is desirable to clarify the bacterial species that indicate the severity of PD available in dogs and develop strategic therapeutic approaches specifically for dogs^34, 62^. These facts underscore the need to investigate with advanced techniques such as shotgun metagenomic and nanopore-based long-read amplicon analysis^63–65^. Also, our study focused intensively on bacteriomes and overlooked the potential role of fungal microbiomes in PD and their impact on bacterial community composition. This area warrants future investigation, especially considering prior research that identified *Cladosporium* spp., *Malassezia restricta*, and *Malassezia arunalokei* as dominant oral fungi^66^.

## Conclusion

This study revealed distinct features in the subgingival plaque microbiome associated with different PD severities. The subgingival bacterial profile in canines changes as PD severity increases, though a process distinct from that observed in humans. Dogs with severe PD exhibited a high prevalence of *Porphyromonas* spp., *Tannerella* spp., and *Treponema* spp. in plaque, pathogens in humans. These results suggest that bacterial composition can indicate the severity of PD. The exacerbation of PD in canines may be attributed to an increase in of highly pathogenic bacteria such as *Porphyromonas* spp. Furthermore, brushing with a gel containing active ingredients was shown to reduce the prevalence of *Porphyromonas* spp. Reducing the prevalence of these pathogenic bacteria with effective materials could be a viable approach for PD control. In conclusion, our findings shed light on the intricate relationship between the subgingival microbiome and PD severity in dogs, highlighting the potential of daily care to control the occupancy of pathogenic microbes. Further research is required to develop an interventional treatment.

## Declaration

### Ethics approval and consent to participate

All procedures were conducted following the Guidelines for Proper Conduct of Animal Experiments of the Science Council of Japan and approved by the KINS Animal Research Ethics Review Committee (approval number: WC003-23-0003). All methods were performed in accordance with the relevant regulations and the ARRIVE 2.0 guidelines.

### Consent for publication

Not applicable

### Data availability statement

The datasets generated and analyzed during the current study are available in the NCBI SRA under the BioProject repository, PRJDB16657.

### Author contributions

MK, YI, and YS designed the experiment. MK, JO, and YS conducted the experimental design. AW and JO conducted the experiments. AW, RN, and KI conducted the data analysis. AW, MK, RN, and KI drafted the manuscript. MK, RN, and KI discussed the results. All authors gave consent to the final manuscript.

## Acknowledgment

The authors thank all the anonymous participants for their contributions: Saori Sumi, Misato Kawamura, Risa Nakamura, and Yasuyoshi Higashide for supporting our sampling; Morgenrot Inc. for providing a computational environment; and Dr. Yoshinori Kawabe for manuscript revisions. We would like to thank Editage [http://www.editage.com] for editing and reviewing this manuscript for the English language. RN is a graduate student of the Medical Innovation Program at Kyoto University and is supported by the JST SPRING program (Grant Number JPMJSP2110).

