## supplementary figures for "Bacterial composition changes in canine plaque over periodontal disease severity and daily care practices"

This file includes 5 supplementary figures.

A

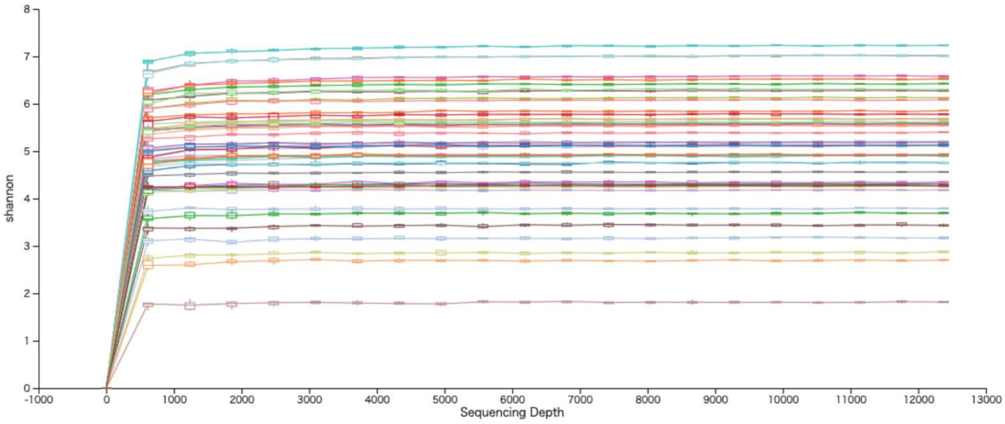


**Supplementary figure 1.** Rarefaction curve showing ASV count per sample depth. (A) Rarefaction curve of the reads used in oral microbiome profiling.

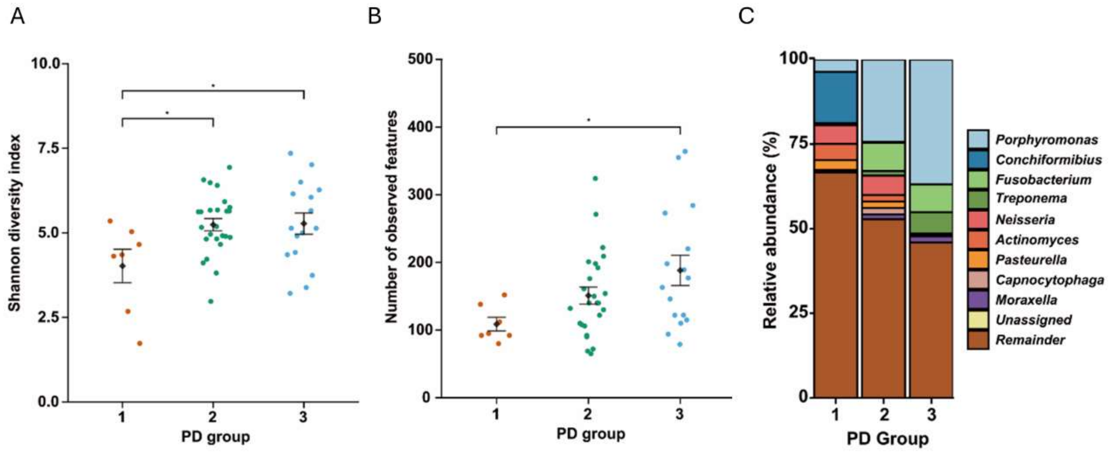


**Supplementary figure 2.** Staked bar chart and Alpha diversity analysis (A) Dot plot of Shannon diversity for each periodontal disease group. (B) Dot plot of observed features for each periodontal disease group. Kruskal–Wallis tests with multiple testing corrections were conducted for both analyzes, and < q = 0.05 is expressed with *. The red dots are Group 1, the green dots are Group 2, and the blue dots are Group 3. (C) Genus-level 100% stacked bar chart. The median relative abundance was calculated for each periodontal disease group. The top 10 genera are indicated, with the others indicated as Remainder (N=48).

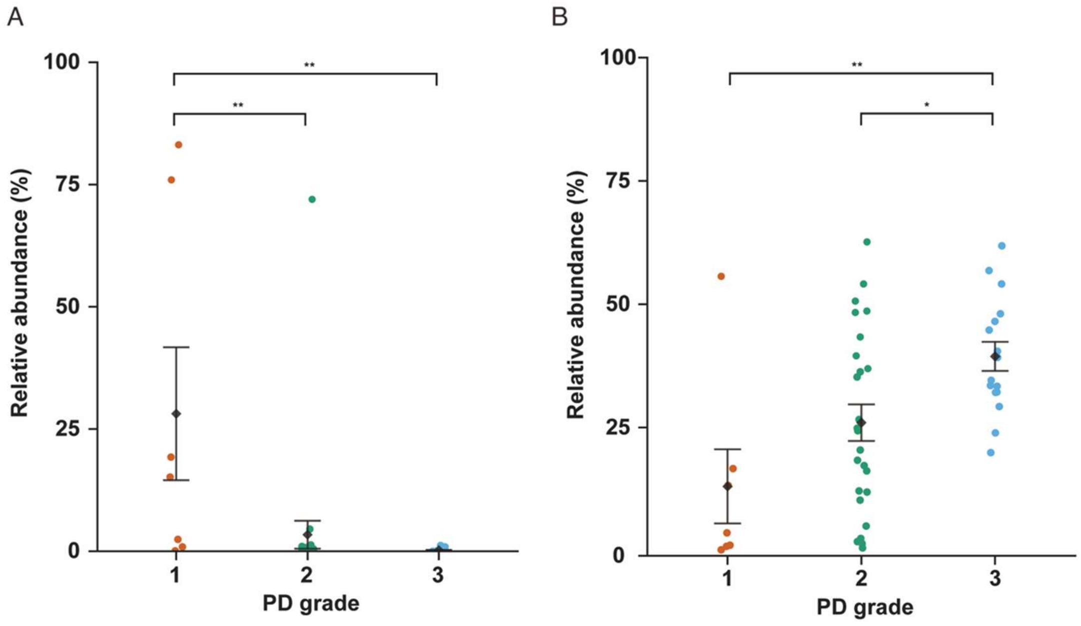


**Supplementary figure 3.** Dot plot of bacteria detected as features. (A) Dot plot of *Conchiformibius*. Its relative abundance is plotted for each periodontal disease group. (B) Dot plot of *Porphyromonas*. Its relative abundance is plotted for each periodontal disease group. Post hoc statistical tests were conducted for both figures, and < p = 0.05 is expressed with *, and < q = 0.01 is expressed with **. The red dots are Group 1, the green dots are Group 2, and the blue dots are Group 3.

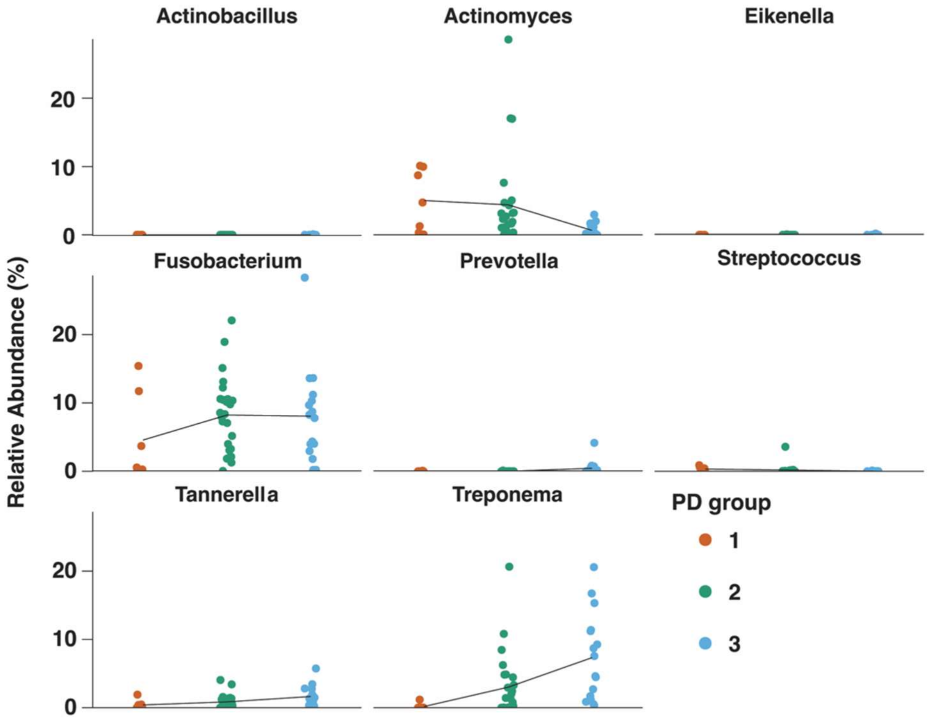


**Supplementary figure 4.** Dot plot of genus-level composition belonging to the Red complex. The relative abundance of each genus is plotted for each periodontal disease group. The red dots are Group 1, the green dots are Group 2, and the blue dots are Group 3.

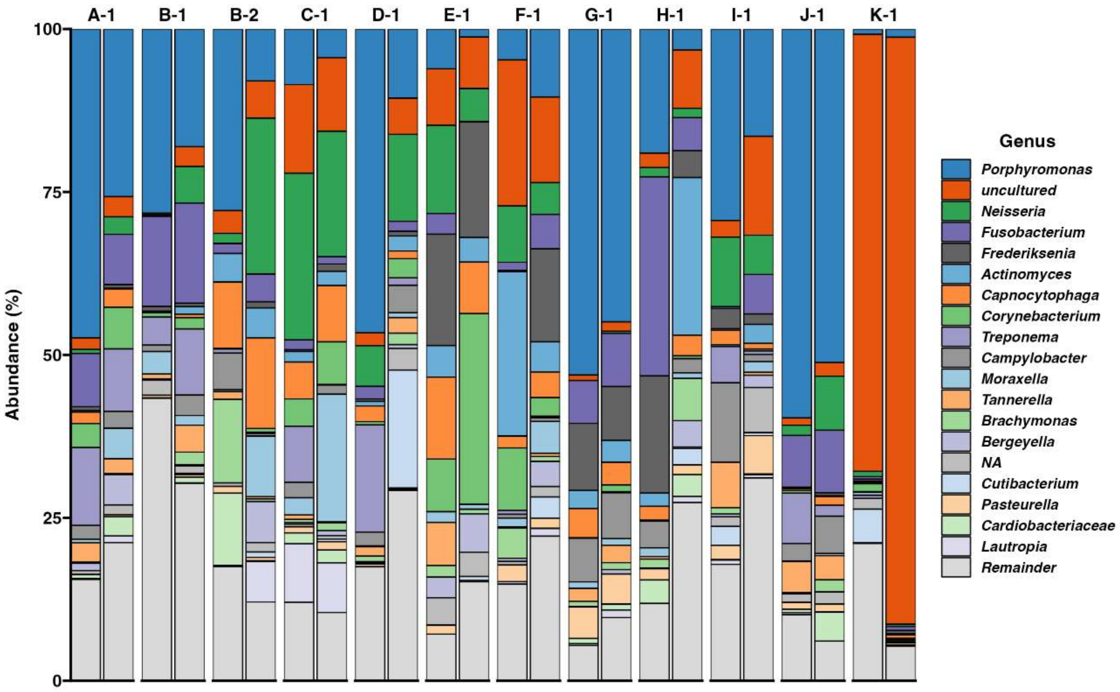


**Supplementary figure 5.** Microbiome analysis before and after toothbrushing with dental gel. The genus-level phylogenetic composition is represented as a 100% stacked bar chart. The top 24 genera are presented by their names, and the rest are grouped as *Remainder*. Two bars per dog represent the relative abundance pre-toothbrushing class (left) and two weeks after the owners' continuous toothbrushing care (right). The alphabet in the subject number indicates the owners of each dog (N=12).
